# Mapping RNA splicing variations in clinically-accessible and non-accessible tissues to facilitate Mendelian disease diagnosis using RNA-seq

**DOI:** 10.1101/727586

**Authors:** Joseph K Aicher, Paul Jewell, Jorge Vaquero-Garcia, Yoseph Barash, Elizabeth J Bhoj

## Abstract

**Purpose:** RNA-seq is a promising approach to improve diagnoses by detecting pathogenic aberrations in RNA splicing that are missed by DNA sequencing. RNA-seq is typically performed on clinically-accessible tissues (CATs) from blood and skin. RNA tissue-specificity makes it difficult to identify aberrations in relevant but non-accessible tissues (non-CATs). We determined how RNA-seq from CATs represent splicing in and across genes and non-CATs.

**Methods:** We quantified RNA splicing in 801 RNA-seq samples from 56 different adult and fetal tissues from GTEx and ArrayExpress. We identified genes and splicing events in each non-CAT and determined when RNA-seq in each CAT would inadequately represent them. We developed an online resource, MAJIQ-CAT, for exploring our analysis for specific genes and tissues.

**Results:** In non-CATs, 39.7% of genes have splicing that is inadequately represented by at least one CAT. 6.2% of genes have splicing inadequately represented by all CATs. A majority (52.8%) of inadequately represented genes are lowly expressed in CATs (TPM < 1), but 6.2% are inadequately represented despite being well expressed (TPM > 10).

**Conclusion:** Many splicing events in non-CATs are inadequately evaluated using RNA-seq from CATs. MAJIQ-CAT allows users to explore which accessible tissues, if any, best represent splicing in genes and tissues of interest.

## Introduction

Exome sequencing is the most advanced standard-of-care genetic test for patients with suspected Mendelian disorders. Yet, the diagnostic rate of exome sequencing is around 31%^1–4^. Genome sequencing, where it has begun being implemented, has been reported to improve upon this diagnostic rate by around 10-15%^1,5,6^. As a result, we are unable to provide a molecular diagnosis for the majority of patients tested with either exome or genome sequencing.

Out of many factors hypothesized to contribute to this diagnostic gap, one particularly significant challenge is our inability to adequately interpret the numerous non-coding and synonymous variants these tests produce^7,8^. These variants can cause disease through various well-described mechanisms but are currently challenging to predict. These difficulties have led to most current clinical pipelines largely ignoring these variants.

One such mechanism by which these variants can cause disease is by altering RNA splicing^9,10^. RNA splicing is the process by which different segments of pre-mRNA are selectively included or excluded and removed as exons and introns to create a mature mRNA (from which proteins are translated) (Figure 1a). This process is highly regulated across developmental stages and tissues and is mediated by the spliceosome and numerous RNA-binding proteins (RBPs) that recognize different conserved sequence elements. Variants in these splice factors can thus alter splicing in *trans*, and intronic and exonic variants can alter splicing in *cis* by changing the strength of existing sequence elements or introducing cryptic ones. These different changes in splicing can alter protein function by inserting or removing part of the mRNA transcript (Figure 1a). Furthermore, they can sometimes cause a frameshift and/or insertion of a premature termination codon, leading to loss of function. Such splicing-altering variants are known to cause Mendelian disorders (i.e. familial dysautonomia, Crouzon syndrome, etc.) and are associated with complex diseases (i.e. Alzheimer’s disease, cancer, etc.)^9,11,12^.

**Figure 1:**
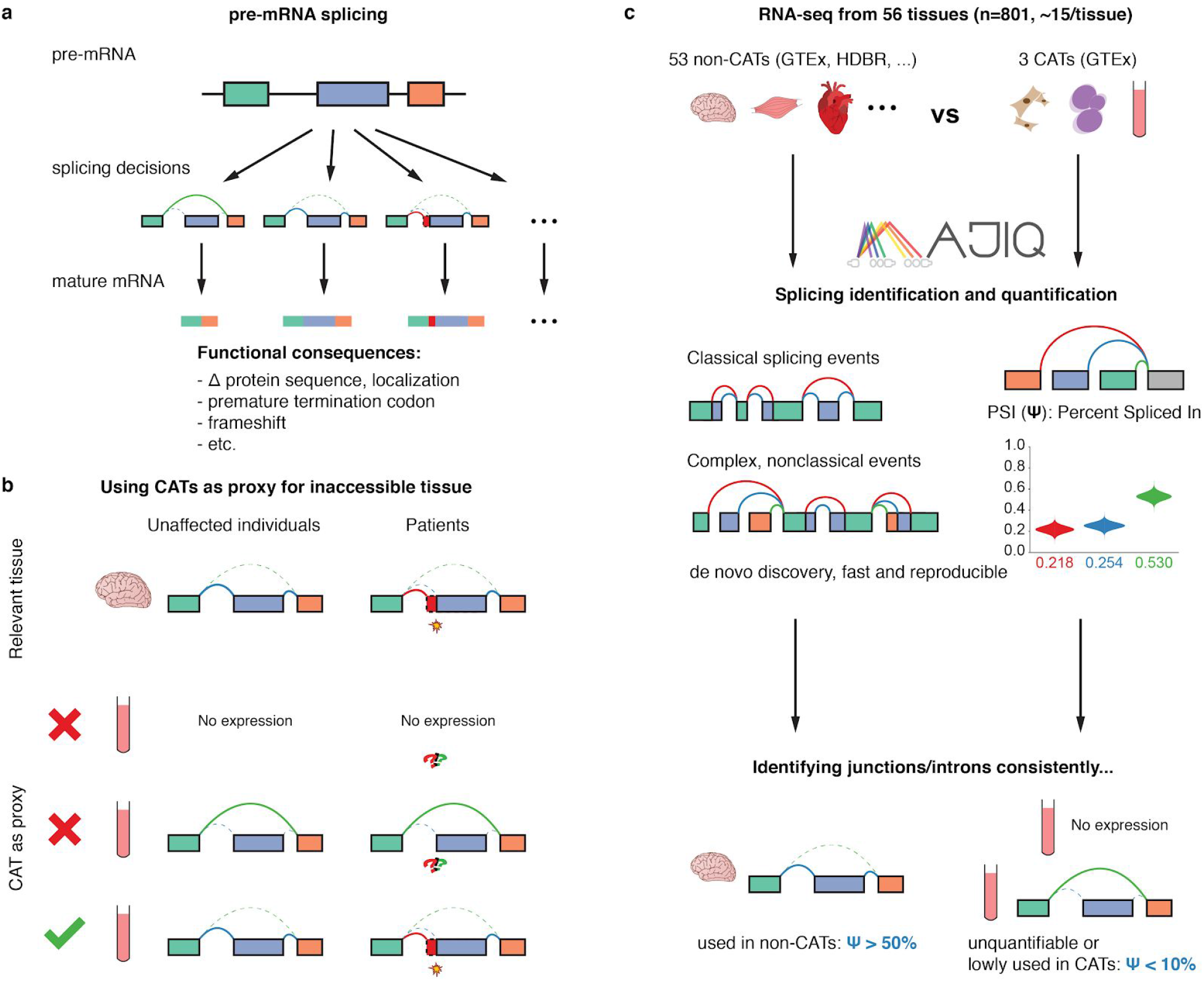
Identification of splicing events inadequately represented by clinically-accessible tissues (CATs). **(a)** Different pre-mRNA splicing decisions can havesignificant, potentiallypathogenic, functionalconsequences. **(b)** Splicing events in inaccessible tissues (non-CATs) can only be adequately represented by accessible tissues as a proxy if the gene is both expressed and similarly spliced. **(c)** We used MAJIQ on RNA-seq samples from 56 different tissues to define and identify inadequately represented splicing events between inaccessible and accessible tissues.

Clinical RNA-seq is one approach by which laboratories can identify splicing aberrations among other changes to the transcriptomes. Previous work in several labs have demonstrated that RNA-seq can enable genetic diagnosis in patients previously unsolved by exome or genome sequencing^13–18^.

As laboratories move to measuring the transcriptome directly with RNA-seq, one challenge they face is tissue specificity. Tissue-specific expression is the most discussed complicating factor of RNA-based analysis, as a gene must be expressed in the tissue to be studied^13,17^. Alternative splicing between tissues is less often addressed, and further complicates analysis. If a tissue other than the tissue of clinical interest is tested, a gene that is expressed in both tissues can still be spliced differently. Thus, splicing defects affecting the tissue of clinical interest might not be realized in the tested tissue despite the gene being expressed in both. Therefore, one tissue can be an adequate proxy for a gene’s splicing in a different tissue only if it is both expressed and spliced similarly (Figure 1b).

Clinicians and researchers can only perform RNA-seq on tissues they have access to. In the clinical setting, these tissues are typically limited to those from blood or skin biopsies: whole blood, EBV-transformed lymphoblasts, and fibroblasts. We refer to these three as <u>**c**</u>linically-<u>**a**</u>ccessible <u>**t**</u>issues (CATs). At the same time, laboratories are often interested in pathology occurring in inaccessible tissues (non-CATs, i.e. brain, heart, etc.).

Several recent studies consider limitations of using RNA-seq from CATs for clinical diagnosis. Frésard et al 2019 demonstrate that RNA-seq in whole blood can make some diagnoses in patients from diverse disease categories^16^. However, Cummings et al 2017, studying a cohort of patients with neuromuscular disease, perform RNA-seq on skeletal muscle biopsies motivated by low gene expression of many known neuromuscular disease genes in whole blood and fibroblasts^13^. Gonorazky et al 2019 further show that they can identify aberrant splicing in muscle that they would not detect from fibroblasts from the same patients^17^. While serving as an important proof of concept of the limitations of RNA-seq with CATs in neuromuscular disease genes, these studies raise the more general question: what are the limitations of RNA-seq in CATs across other non-CATs and genes in general, and how can one quantify them? An answer to this question could inform the selection of the best clinically-accessible tissue (if any) to send for RNA-seq for different patients in clinical practice by evaluating the degree to which each CAT is able to faithfully represent splicing in genes and tissues of interest for a given patient’s phenotype.

We address the question by considering splicing in non-accessible vs accessible tissues in terms of splicing events. We model splicing events as local splicing choices either starting or ending at a single exon (the reference exon) in a given gene. These local choices are between splice junctions or intron retention between exons that are included in the gene’s transcripts. These splicing events include constitutive splice junctions and local splicing variations (LSVs)^19^, which are splicing events where the reference exons can be spliced to multiple RNA segments, thus allowing variation.

We identify and quantify splicing events from RNA-seq data using the recently updated MAJIQ 2.1 toolkit for splicing detection, quantification, and visualization^19^. Previous studies have shown MAJIQ is able to accurately detect and quantify splicing variations, comparing favorably to other tools and offering unique advantages for the specific task we consider here^20,21^. Briefly, MAJIQ can robustly detect LSVs by combining given transcriptome annotations with *de novo* (unannotated) splice junctions and retained introns found in the input RNA-seq data in order to detect novel exons and LSVs. For each LSV junction (or retained intron), MAJIQ quantifies its relative inclusion compared to the other LSV junctions, measured as percent splicing inclusion (PSI or Ψ ∈ [0, 100]), in any given sample, allowing us to quantify and compare splicing between samples and tissue-types. Previous work has shown that MAJIQ highly correlates with RT-PCR validation experiments (*r=0.97*), the gold standard in the RNA field, and identifies differentially spliced events between RNA-seq samples with high reproducibility (*RR=78%*)^21^.

When using splicing in CATs as a proxy to splicing in some other tissue of interest, we consider three possible scenarios or splicing event categories (Figure 1b): (1) the event is unquantifiable in the CAT due to low gene expression and/or sequencing depth, (2) the event is quantifiable but spliced differently, and (3) the event is quantifiable and not spliced differently. We further focus on splicing events that are consistently included, meaning that they are similarly quantified in nearly all samples for a given tissue-type. Naturally, categorizing events into these scenarios depend on the thresholds used to define them. Here, we define consistently spliced events in non-CATs to be events with a junction or retained intron with Ψ > 50% in more than 85% of samples (Figure 1c). We emphasize finding the subset of these events that correspond with either of the first two scenarios where splicing in a CAT inadequately represents splicing in the non-CAT. We define these events as those that are unquantifiable or have Ψ < 10% in more than 85% of a CAT’s samples.

In this work, we analyze 53 adult tissues in GTEx^22^ and 3 fetal tissues from HDBR^23^ (cerebellum, cortex) and ArrayExpress accession E-MTAB-7031^24^ (heart) (Figure 1c). We map all transcriptome variations across these datasets, contrasting splicing between CATs and non-CATs. We make our analyses accessible as an online resource which we call MAJIQ-CAT: <https://tools.biociphers.org/majiq-cat>. This online resource has been designed for clinicians and researchers interested in obtaining patient RNA-seq in the context of Mendelian disease. With MAJIQ-CAT, these users can explore how faithfully different CATs represent splicing in their specific genes and tissues of interest, informing their choice of patient tissue to collect. Finally, we discuss implications for RNA-seq in clinical practice and the need for alternative solutions for the genes and tissues that are inadequately represented by CATs.

## Materials and Methods

### Sample selection criteria

We used RNA-seq data for samples from 56 different tissue-types: 53 adult tissues and 3 fetal tissues. We obtained samples for all 53 adult tissue-types from the Genotype-Tissue Expression project (GTEx; dbGaP accession phs000424). Meanwhile, we obtained samples for fetal cerebral cortex and cerebellum from the Human Developmental Biology Resource (HDBR; ArrayExpress accession E-MTAB-4840), and we obtained samples for fetal heart from ArrayExpress accession E-MTAB-7031.

We restricted sample selection from each of these datasets/tissue-types using available metadata as follows. We restricted selection to unique donors per tissue-type. This restriction was relevant to both GTEx and HDBR, which include donors contributing multiple sample per tissue-type. Available HDBR metadata did not suggest criteria for preferring one sample over another, so we restricted selection to the first available sample per donor. However, GTEx metadata includes information on the number of megabases per sample, so we restricted selection to the sample with the largest size. GTEx metadata also included further information that we used; specifically, we further restricted selection to samples that: (1) were hosted by NCBI, (2) had matched whole-genome sequencing data, (3) had an average spot length of 152bp, (4) were not flagged by GTEx as a sample to remove (SMTORMVE), and (5) had a RIN score greater than 6.

Given these restrictions, we selected up to 15 samples per tissue-type for further analysis. We chose 15 samples as the maximum number of samples per tissue group for because preliminary analysis using MAJIQ indicated that reproducibility for tissue-specific differential splicing analysis saturates with around 15 samples in GTEx (data not shown). Consequently, when there were more than 15 samples meeting the above criteria for a particular tissue-type, we randomly selected 15 samples among them. For the other tissue-types, we kept all samples meeting criteria for further analysis.

### Sample read alignment

We aligned RNA-seq reads from the selected samples to the human genome for splicing analysis with MAJIQ using the following procedure. We downloaded selected samples as FASTQ files using SRA Tools (v2.9.6)^25^. We performed quality and adapter trimming on each sample using TrimGalore (v0.4.5)^26^. We used STAR (v2.5.3a)^27^ to perform a two-step gapped alignment of the trimmed reads to the GRCh38 primary assembly with annotations from Ensembl release 94^28^.

### Gene expression quantification

We quantified gene expression in each of the samples. We quantified transcript abundances in TPM for using Salmon (v0.13.1)^29^ quasi-mapping on the trimmed reads for each sample with a transcriptome built from Ensembl release 94 annotations. We aggregated the quantifications to gene expression by taking the sum of abundances for the transcripts associated with each gene.

### Splicing identification and quantification using MAJIQ

First, we used MAJIQ (v2.1)^19^ with Ensembl release 94 annotations to identify/model the set of all possible splicing events across our samples. We then quantified these splicing events for each sample, considering an event to be quantifiable for a given sample if it had at least one junction with at least 10 supporting reads starting from at least 3 unique positions. We estimated the percent spliced in (PSI or Ψ) for the junctions and retained introns in each quantifiable splicing event. For the quantifiable LSVs, we used MAJIQ PSI to estimate PSI. Meanwhile, we assigned Ψ = 100% to the quantifiable constitutive junctions, as they were the only choice for inclusion in their respective events.

Finally, we identified and filtered out ambiguous splicing events per sample. We defined ambiguous splicing events as events containing junctions or retained introns that were in quantified splicing events assigned to more than one gene. These ambiguous assignments occur because of the presence of overlapping genes, especially combined with the unstranded nature of the RNA-seq experiments we used. The resulting nonambiguous quantifications were used to identify relevant consistent and tissue-specific differences between CATs and non-CATs.

### Identifying relevant splicing events

In order to determine the extent to which splicing in non-CATs is inadequately represented by splicing in CATs, we first defined which splicing events for each non-CAT we would consider changes in usage for. For each non-CAT, we consider the set of consistent splicing events, which are splicing events with a junction or retained intron that is highly included in nearly all samples for their tissue-type. Specifically, we considered splicing events with a junction or retained intron quantified as Ψ > 50% in more than 85% of the samples for each non-CAT.

We then evaluated how well splicing quantified in CATs reflected splicing in these consistent splicing events. To do so, we identified the subset of these events for which usage in CATs was consistently low or unquantified. Specifically, we identified which events were either unquantifiable or had Ψ < 10% for the same junction or retained intron in more than 85% of the samples for each CAT. We call these splicing events inadequately represented in their respective CAT.

### Analysis of genes with consistently used splicing events

We then aggregated information about these consistent and inadequately represented splicing events to their respective genes for each tissue. That is, we determined which genes had consistent splicing events in each non-CAT and the subset of these genes for which these events were inadequately represented for each CAT. We evaluated gene expression for the inadequately represented genes to assess how inadequately represented splicing related to low gene expression vs tissue-specific alternative splicing. We also evaluated which of the inadequately represented genes were annotated as disease-causing. We obtained our list of disease-causing genes by combining annotations from ClinVar^30^ and HGMD 2018.3^31^.

### Data access and software

Sequencing data used for this analysis are available in dbGaP under accession phs000424 and ArrayExpress under accessions E-MTAB-4840 and E-MTAB-7031. Software versions, resources, and specific parameters used are listed in Table S1. The analysis was implemented for reproducible execution as a Snakemake pipeline^32^. Links to source code for the analysis and online resource, MAJIQ-CAT, are listed in Table S2.

## Results

Our sample-selection procedure yielded a dataset with *n*=801 RNA-seq samples for 53 non-CATs and 3 CATs (Table S3). 762 samples came from GTEx, 30 fetal brain samples came from HDBR, and 9 fetal heart samples came from E-MTAB-7031. We selected and processed 15 samples for each tissue except for bladder, cervix (ectocervix and endocervix), fetal heart, and fallopian tube, where we selected all available samples that met our criteria.

Across all samples, we identified a total of 227,909 quantifiable splicing events (129,582 LSVs and 98,327 constitutive junctions) in 25,732 genes. Per sample, we quantified a median of 108,050 splicing events (67,556 LSVs and 40,701 constitutive junctions) in 12,805 genes. We then identified and removed ambiguous splicing events with junctions or retained introns associated with multiple genes, leaving a total of 212,855 splicing events (122,015 LSVs and 90,840 constitutive junctions) in 24,926 genes with a per-sample median of 99,991 splicing events (62,364 LSVs and 37,742 constitutive junctions) in 12,118 genes.

Among quantified LSVs and constitutive junctions, we identified in each non-CAT a median of 67,486 junctions or retained introns in 10,043 genes that were consistently used (Figures 2a, S1; Table S4). Looking at these same events in CATs, we found that 28.1% were inadequately represented in at least one CAT (3,885 or 39.7% of genes) (Table S5). 4.6% were inadequately represented by all CATs (609 or 6.2% of genes).

**Figure 2:**
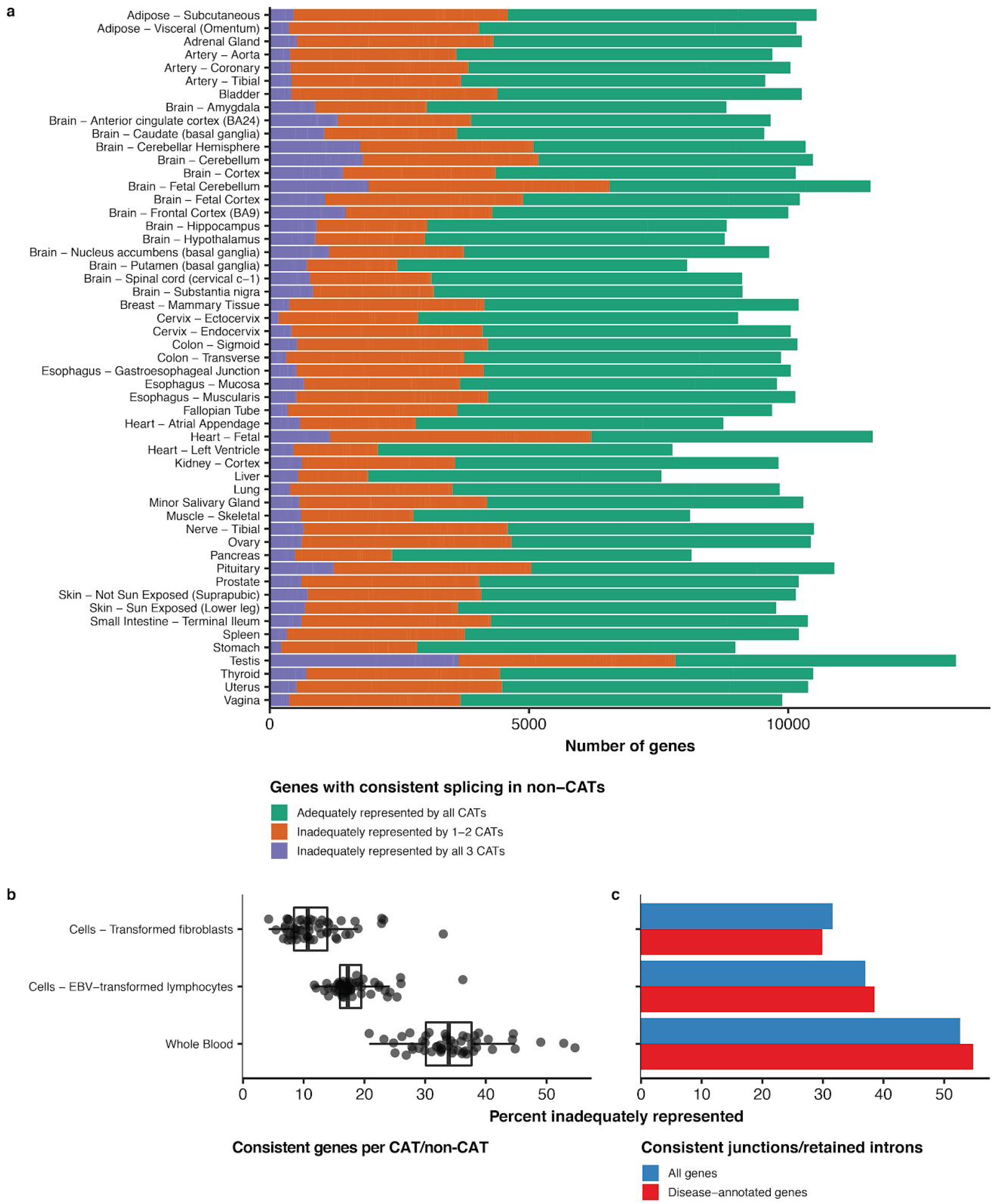
Mapping transcriptome variations identified in CATs vs non-CATs. **(a)** Of an average of 10,043 genes with consistently spliced events per non-CAT, 3,885 (39.7%) were inadequately represented in at least oneCAT, with 609 (6.2%) being inadequately represented by all CATs. **(b)** The percentages of inadequately represented genes over non-CATs was lowest in fibroblasts and highest in whole blood. **(c)** The percentage of junctions/retained introns that were consistently used in at least one non-CAT that were inadequately represented by each CAT was lowest in fibroblasts and highest in whole blood.

We compared the quantities of inadequately represented splicing per CAT and non-CAT. The median percentage of inadequately represented genes across non-CATs was 10.7% for fibroblasts in comparison to 17.3% for EBV-transformed lymphoblasts and 33.9% for whole blood (Figure 2b). The percentage of inadequately represented genes was lowest in fibroblasts and highest in whole blood for each non-CAT except for spleen, for which the percentage was lowest in EBV-transformed lymphoblasts (Figure S2). Considering all consistently spliced junctions/retained introns across non-CATs, we found that the percentage of inadequately represented splicing was also lowest in fibroblasts and highest in whole blood (Figure 2c).

We further investigated the expression and pathogenicity of inadequately represented genes. The maximum median gene expression across inadequately representing CATs was less than 1TPM in 52.8% of inadequately represented genes (Figure 3a). However, an average of 5.8% of inadequately represented genes per non-CAT (227 genes) are expressed with greater than 10TPM. Meanwhile, a median of 29.2% of inadequately represented genes per non-CAT were annotated as disease-causing in either ClinVar or HGMD (Figure 3b).

**Figure 3:**
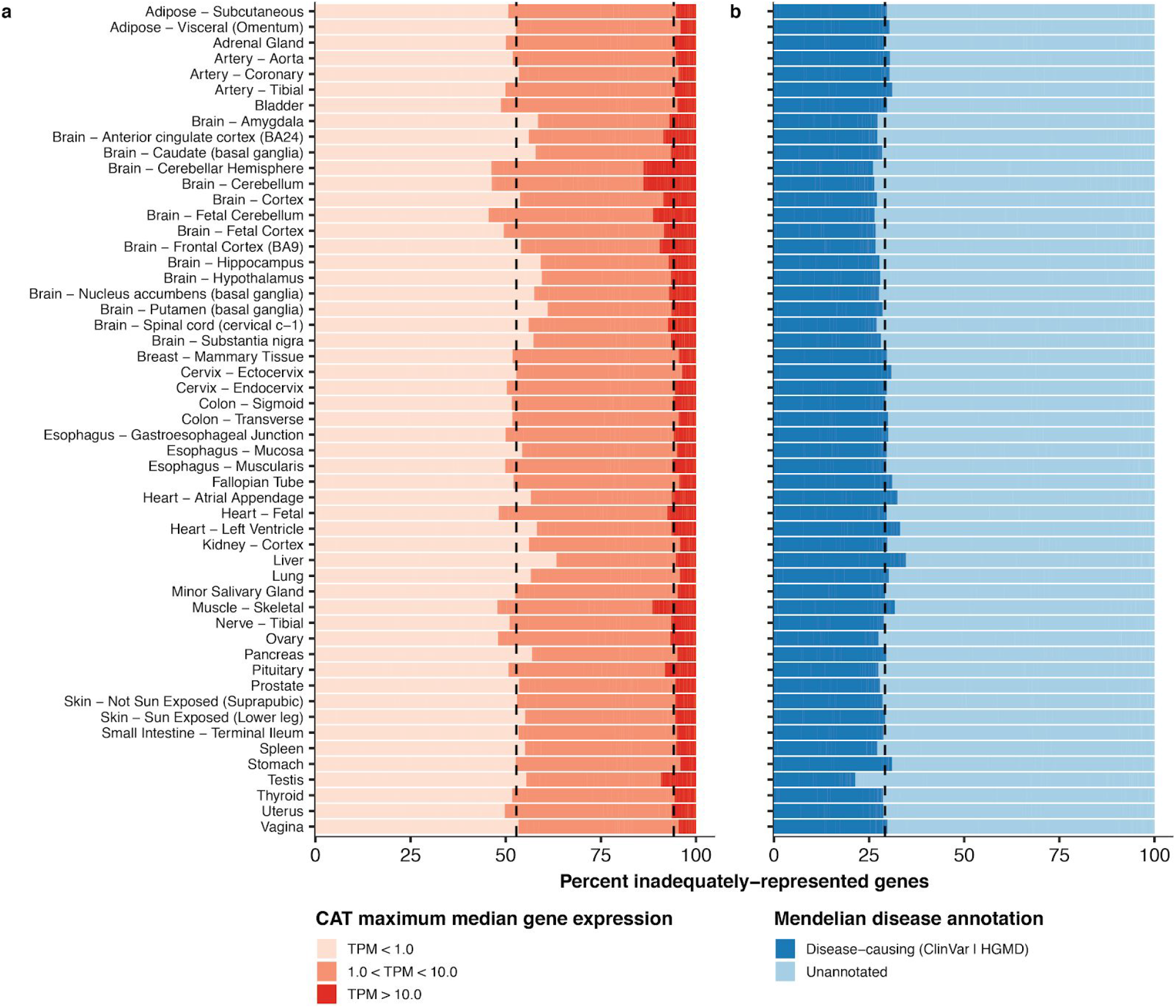
Expression and pathogenicity of inadequately represented genes. **(a)** The majority of inadequately represented genes are lowly expressed (TPM < 1) inCATs, but an average of 227 genes (5.8%) are well expressed (TPM > 10) in at least one inadequately representing CAT. **(b)** An average of 29.2% of inadequately represented genes are annotated as disease-causing.

To facilitate the interrogation of specific genes and splicing variations of interest by clinicians, we developed MAJIQ-CAT <https://tools.biociphers.org/majiq-cat>. MAJIQ-CAT is an online resource that provides panels with which users can select genes and non-CATs to look at how well CATs represent splicing across their genes of interest, both globally (Figure 4a) and looking at individual splicing events in a specific gene (Figure 4b). Genes can be selected from predefined lists of genes (i.e. from ClinVar, ClinGen, etc.) or custom lists provided by the user either interactively or by uploading a text file. Non-CATs can be selected similarly. Changes to these inputs automatically regenerate plots and tables describing the consistent and inadequately represented genes. Individual genes can further be explored by clicking their names to load an additional page which displays their tissue-specific gene expression and splicing events. For example, if a laboratory was interested in studying intellectual disability as a phenotype, they could focus the brain non-CATs and genes associated with the corresponding HPO term (HP:0001249), finding 1,229 genes with consistent splicing in at least one of the brain tissues (Figure 4a). They could further look for genes that are expressed but inadequately represented by filtering the table by expression; in this example, setting a minimum of TPM > 100 yields a list of 18 genes. Clicking into one of the resulting genes (i.e. *MEF2C*) leads to another page with comparisons of splicing in the CATs and the brain tissues, demonstrating where the inadequately represented splicing events are and the distributions of PSI in each tissue for each event (Figure 4b).

**Figure 4:**
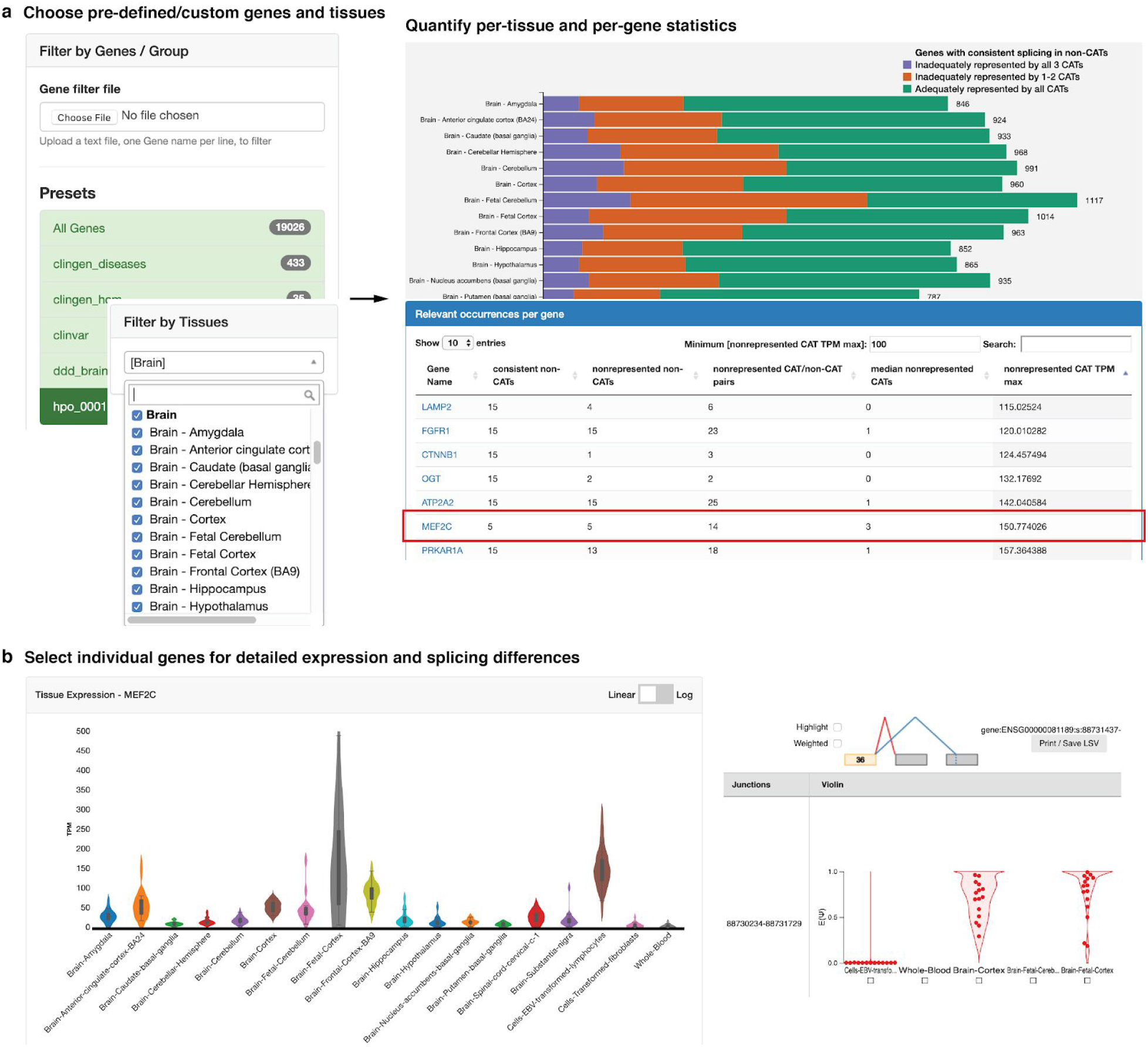
MAJIQ-CAT enables clinicians and scientists to explore inadequate representation of splicing by CATs in specific genes and tissues of interest. **(a)** MAJIQ-CAT allows users to choose from predefined or custom gene sets and tissues (left) to quantify and understand the user-specific relevant limitations of RNA-seq in different accessible tissues (right). **(b)** Users can further explore individual genes for tissue-specific differences in gene expression and splicing. Shown here is a closer look at the gene *MEF2C*, with a violin plot of its expression in CATs and selected non-CATs (left) and violin plots of PSI for one of its inadequately represented splicing events (right). See main text for more details.

## Discussion

In this study, we present a comprehensive analysis of RNA splicing events that consistently take place in clinically-inaccessible tissues, focusing on how corresponding events take place in clinically-accessible tissues. While clinicians and scientists are often interested in what takes place in the inaccessible tissues as part of disease pathology, laboratories can only measure the accessible tissues to estimate by proxy. Thus, these results inform clinicians and scientists as to where RNA-seq is limited, especially with respect to previously under-appreciated tissue-specific splicing, and suggest when specific clinically-accessible tissues should be preferred over others or when alternative approaches to clinical RNA-seq are needed. By making the results interactively accessible through MAJIQ-CAT, we enable clinicians and scientists to more directly explore how these limitations impact specific genes and tissues of interest to them.

Previous studies analyzing patient RNA-seq from CATs demonstrated that these data could be used to identify rare disease genes and variants from a variety of disease categories^14,16^. These studies demonstrate that although clinicians are typically limited to using CATs for patient RNA-seq, RNA-seq in those tissues can still improve the molecular diagnostic rate for suspected Mendelian disorders by identifying changes that were not identified using exome or genome sequencing alone. Our study provides an orthogonal, but related, result. Because we are typically limited to using CATs for patient RNA-seq, there are splicing events found in disease-relevant tissues and genes that will consistently be a blind spot in such studies.

Our study found that 39.7% of genes with consistent splicing events per non-CAT are inadequately represented by at least one CAT. This implies that clinicians and scientists interested in how one of the inadequately-represented genes are spliced in the non-CAT in patients need to be careful about which clinically-accessible tissues they measure as a proxy because at least one of the accessible tissues will not represent the splicing events well. We show that many of these genes are considered disease-causing (29.2% of the inadequately represented genes); thus, understanding these limitations are increasingly clinically relevant as RNA-seq enters clinical practice.

Considering these 39.7% of genes with inadequately represented splicing, the majority (52.8%) were associated with low gene expression (TPM < 1), as expected. However, we still find that 227 genes per non-CAT are highly expressed (TPM > 10) but spliced differently in CATs. The limitations of these genes for clinical RNA-seq would be missed by previous expression-first analyses, highlighting the novelty and impact of our splicing-first analysis.

For the other 60.3% of genes, we note that splicing in CATs may still not always adequately represent splicing in non-CATs. While they may not pass the stringent thresholds we set to define inadequately represented splicing present in most samples for a CAT (Ψ < 10% or unquantifiable in more than 85% of samples), splicing inclusion may take intermediate values or be highly variable between samples. Furthermore, even for splicing variations that are similar between tissues, they may still involve different tissue-specific regulation by different tissue-specific factors. Thus, while we might expect variants in tissue-independent splicing sequence elements (e.g. canonical splice sites) to impact the different tissues similarly, variants in tissue-specific splicing enhancers or silencers could lead to tissue-specific defects that would not be represented by CATs.

One important conclusion from the analysis performed here is that for the 3,276 genes per non-CAT that are inadequately represented by one or two CATs, at least one CAT offers a better representation of the gene’s splicing than the others. Thus, our study implies that researchers interested in one of these genes and tissues should have a preference for which clinically-accessible tissue to collect. Summarizing across all genes with consistent splicing, we found that fibroblasts almost always had the lowest percentage of inadequately represented genes. Thus, our results suggest that researchers interested in all genes and tissues equally should prefer collecting patient fibroblasts if possible. However, clinicians and scientists are often interested in specific genes or tissues relevant to a specific biological process. Our online resource, MAJIQ-CAT, will enable clinicians, scientists, and laboratories to interactively explore which CATs are most relevant for representing the biology they care about and which genes and splicing events are most affected.

Another important conclusion of this study is that there are 609 genes per non-CAT that are inadequately represented by all CATs. For these genes, using RNA-seq in any CAT as a proxy would have many limitations for studying splicing. In these cases, alternative approaches are likely necessary. One possible path forward is the use of *in vitro* differentiation/transdifferentiation of CRISPR-iPSCs/patient-derived cells towards tissue-types of interest. Gonorazky et al 2019 illustrated this possibility for transdifferentiated myotubes from patient fibroblasts as an alternative to skeletal muscle biopsy, although how these results would translate to other, more inaccessible, tissues remains to be explored^17^. Another possible path forward is the use of *in silico* models of splicing^33–39^. Previous work in several labs have developed models of tissue-specific splicing but do not directly train models on genetic variants^33–37^. Recent work by Cheng et al 2019 directly trains models to predict splicing changes using genetic variants but does not account for tissue-specificity^39^. Future developments combining aspects of these models to produce predictions of tissue-specific splicing as a consequence of genetic variants could help us understand potential splicing defects in those genes where we do not have a good proxy.

In summary, in this study, we demonstrated and quantified the limitations of CATs to serve as a proxy for non-CATs for RNA splicing measured by RNA-seq. We highlighted how alternative splicing contributes to these limitations in addition to tissue-specific gene expression. In addition, we developed and have made available an online resource, MAJIQ-CAT, that will allow clinicians and scientists to directly explore how these limitations affect specific genes and tissues of interest. MAJIQ-CAT will be of particular use for determining tissues to study for genes that are only inadequately represented in some but not all CATs. For the genes inadequately represented by all CATs, future work on alternative approaches to estimate splicing defects in patients will be necessary to improve clinical diagnoses.

## Acknowledgements

Research reported in this publication was supported by the National Institute of General Medical Sciences of the National Institutes of Health under award number R01GM128096. The content is solely the responsibility of the authors and does not necessarily represent the official views of the National Institutes of Health. J.K.A. acknowledges salary support by NIH/NICHD fellowship F30HD098803.

## Figure Legends

**Figure S1:**
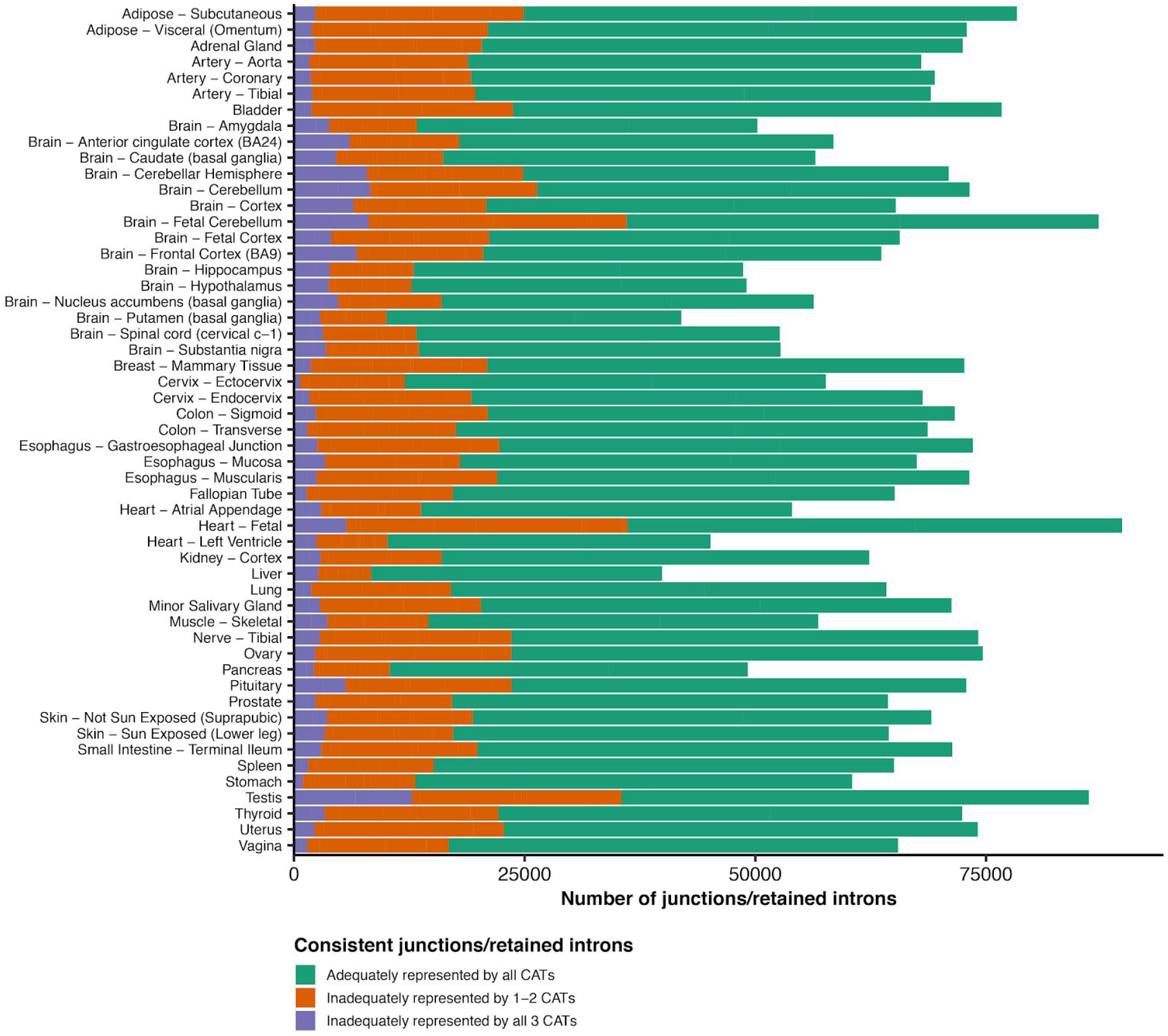
Mapping transcriptome variations identified in CATs vs non-CATs. Of an average of 67,486 consistently spliced junctions or retained introns per non-CAT, 19,213 (28.1%) were inadequately represented in at least one CAT, with 2,828 (4.6%) being inadequately represented by all CATs.

**Figure S2:**
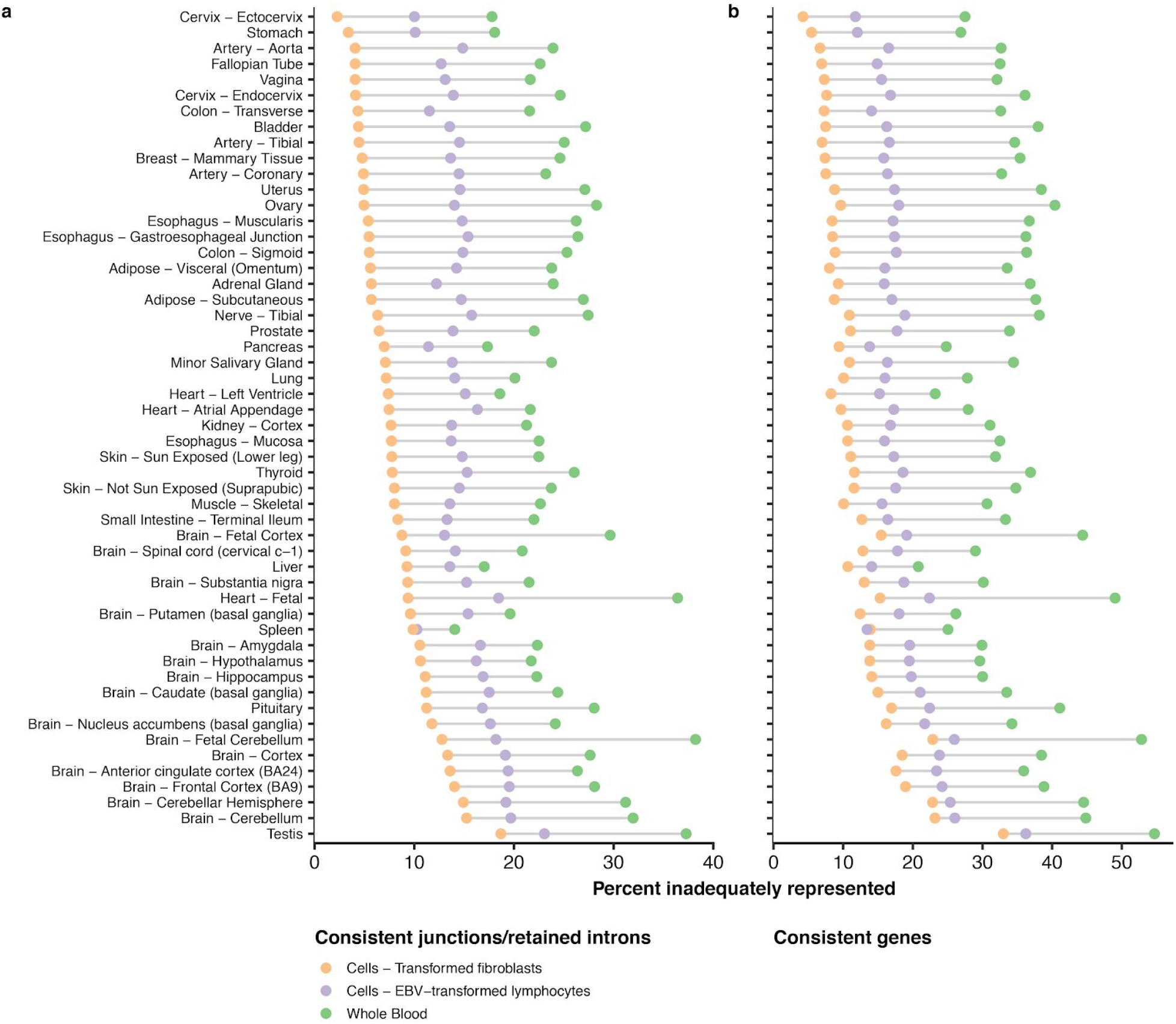
Inadequately represented splicing per CAT and non-CAT. **(a)** Considering all consistent splicing events per non-CAT, the percentage of inadequately represented junctions/retained introns per CAT was always lowest for fibroblasts and highest for whole blood. **(b)** Considering all genes with consistent splicing events per non-CAT, the percentage of genes with inadequately represented splicing was lowest for fibroblasts and highest for whole blood for all non-CATs except spleen.

