## Supplementary figures and images for "Mapping RNA splicing variations in clinically-accessible and non-accessible tissues to facilitate Mendelian disease diagnosis using RNA-seq"

### figure_s1.pdf

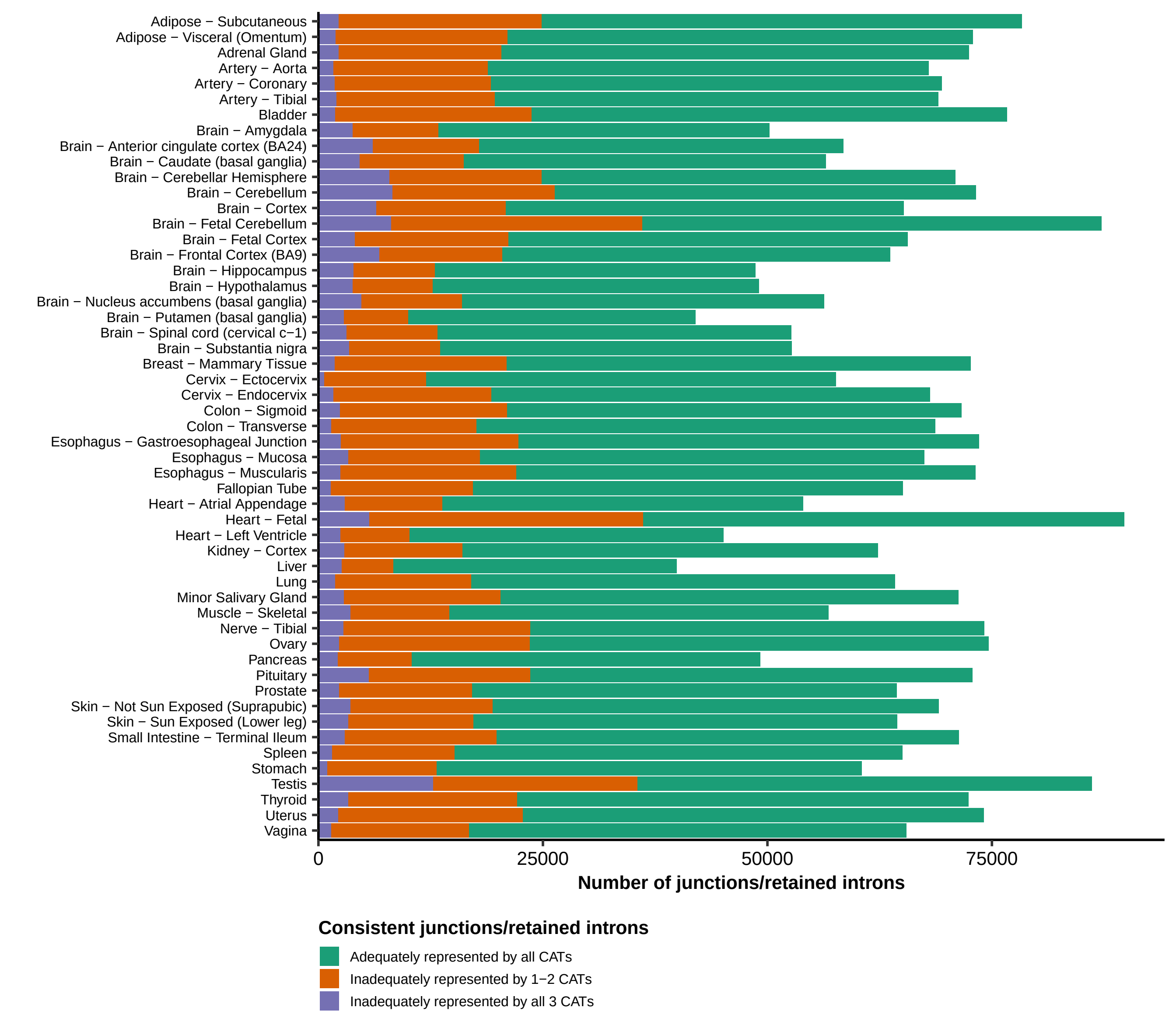

### figure_s2.pdf

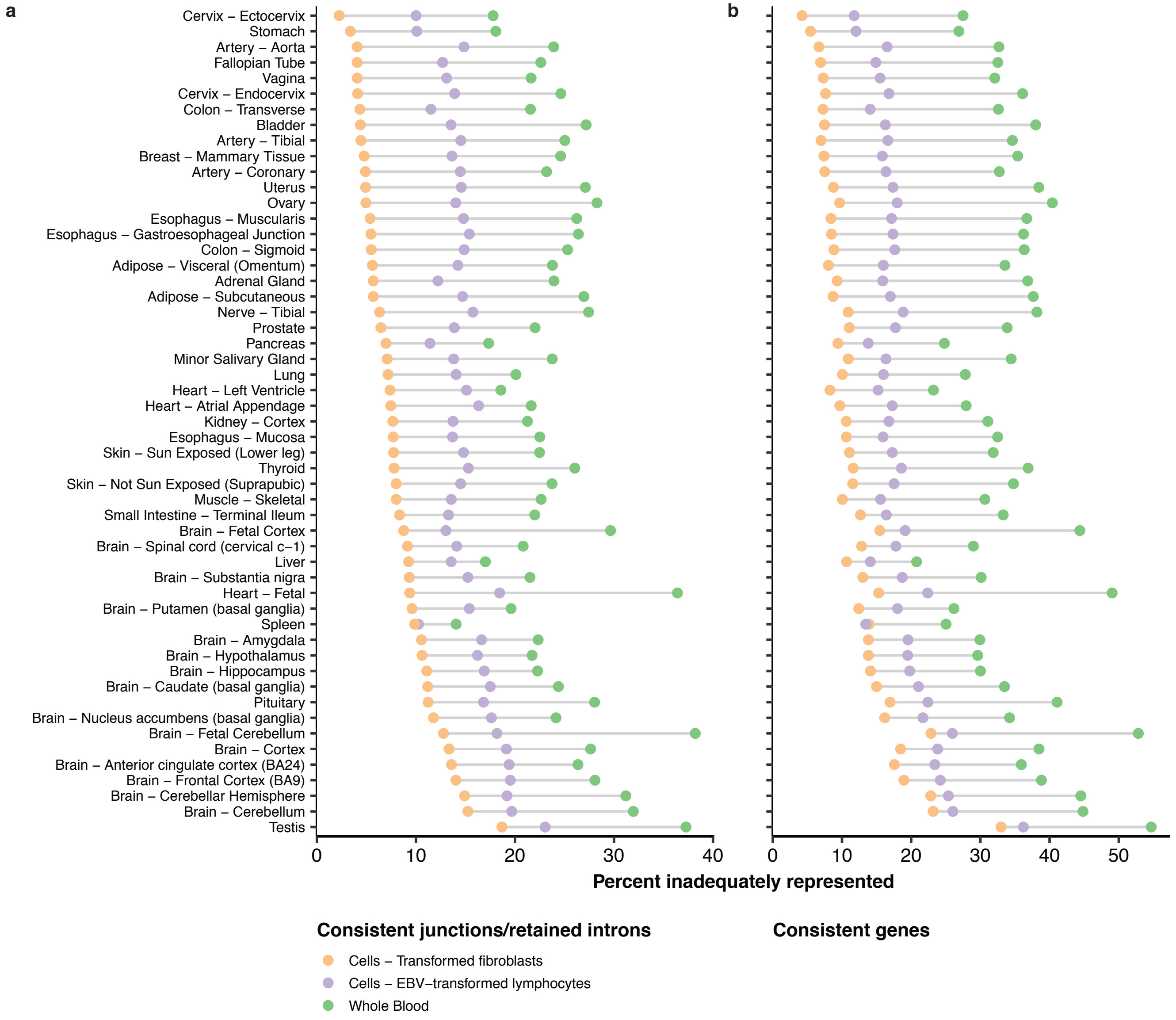
